# Integrating Genomic Correlation Structure Improves Copy Number Variations Detection

**DOI:** 10.1101/2020.04.08.032680

**Authors:** Xizhi Luo, Fei Qin, Guoshuai Cai, Feifei Xiao

## Abstract

Copy number variation plays important roles in human complex diseases. The detection of copy number variants (CNVs) is identifying mean shift in genetic intensities to locate chromosomal breakpoints, the step of which is referred to as chromosomal segmentation. Many segmentation algorithms have been developed with a strong assumption of independent observations in the genetic loci, and they assume each locus has an equal chance to be a breakpoint (i.e., boundary of CNVs). However, this assumption is violated in the genetics perspective due to the existence of correlation among genomic positions such as linkage disequilibrium (LD). Our study showed that the LD structure is related to the location distribution of CNVs which indeed presents a non-random pattern on the genome. To generate more accurate CNVs, we therefore proposed a novel algorithm, LDcnv, that models the CNV data with its biological characteristics relating to genetic correlation (i.e., LD). To evaluate the performance of LDcnv, we conducted extensive simulations and analyzed large-scale HapMap datasets. We showed that LDcnv presents high accuracy, stability and robustness in CNV detection and higher precision in detecting short CNVs compared to existing methods. We also theoretically demonstrated the correlation structure of CNV data, which further supports the necessity of integrating biological structure in statistical methods for CNV detection. This new segmentation algorithm has a wide scope of application with next-generation sequencing data analysis and single-cell sequencing analysis.

**Author Summary:** Copy number variants (CNVs) refers to gains or losses of the DNA segments in comparison to a reference genome. CNVs have garnered extensive interests in recent years as they play an important role susceptibility to disorders and diseases such as autism, schizophrenia and cancer [1-7]. Although innovation in modern technology is promoting the discoveries related to CNVs, the methodology for CNV detection is still lagging, which limits the novel discoveries regarding the role of CNVs in complex diseases. In this study, we are proposing a novel segmentation algorithm, LDcnv, to accurately locate the breakpoints or boundaries of CNVs in the human genome. Instead of utilizing an independent assumption of the signal intensities as has been used in traditional segmentation algorithms, LDcnv models the correlation structure in the genome in a change-point CNV detection model, which allows for accurate and fast computation with a whole genome scan. Our study showed strong theoretical evidence of the existence of correlation structure in real CNV data, and we believe that taking this evidence into consideration will improve the power of CNV detection. Extensive simulation studies have demonstrated the advantage of the LDcnv algorithm in stability, robustness and accuracy over existing methods. We also used high-quality CNV profiles to further support the superior performance of the LDcnv algorithm over existing methods. The development of the LDcnv algorithm provides great insights for new directions in developing CNV detection tools.

## 1. Introduction

Copy number variants (CNVs), as a major source of genetic variation in the human genome, are gains or losses of the DNA segments in comparison to a reference genome. Recently, copy number variation has garnered considerable interest as it plays an important role in the susceptibility to disorders and diseases such as autism, schizophrenia and cancer [1-7]. It has been found that approximately 12% of the genome is subject to CNVs and nearly 80% of cancer genes harbor CNVs [8]. To date, CNV studies have been intensively conducted in many disease types which demonstrates that CNVs account for an abundance of genetic variation and play essential roles in the etiology of cancer [9-11], autoimmune diseases [12, 13] and neurological diseases [14, 15]. Specifically, about 200 CNVs have been found to be associated with breast cancer risk, among which 21 had prognostic potential [9]. Also, copy number gain of beta-defensin genes has been revealed to be associated with increased risk of psoriasis in three independent cohorts of European origin [9, 14, 16]. Two recent reports also illustrated the possible roles of CNVs in lung cancer predisposition [17, 18]. In one of these two reports, an approximately 2-fold increased risk was *observed* among carriers with deletion of the gene coding region of *WWOX* compared to non-carriers [17].

Although the potential clinical application of CNVs still remains uncertain, understanding the mechanisms underlying these influences will be instrumental for many basic research areas. Consequently, the detection and association of CNVs with quantitative traits and clinical phenotypes comprise critical steps toward a better understanding of disease etiology. However, due to the complexity of CNV genetics as well as numerous factors in the data generation and computational analyses that may lead to spurious associations, the discovery of CNVs in human diseases is still inadequate, which places obstacles in the path of utilizing CNVs as important biomarkers for clinical applications.

Technically, the detection of CNVs is the finding of breakpoints or boundaries of copy number regions from the genotyping signals, the step of which is called chromosomal segmentation. Change-point tests have been commonly used and implemented in many software and tools for chromosomal segmentation [19-21]. Among them, circular binary segmentation (CBS) is widely used and is based on an exhaustive test [22]. This segmentation algorithm has been widely utilized in whole exome sequencing data tools such as ExomeCNV [23] and CODEX [24]. In EXCAVATOR [25], a shifting level model segmentation algorithm was used to incorporate the distance between two genetic sites. More recently, a novel segmentation procedure was utilized in modSaRa that adopted a local search strategy and was demonstrated to be suitable for whole genome analysis with low computational complexity [26-28]. Nevertheless, all of these algorithms were developed with a strong assumption of independent observations in the genetic loci and they assume each locus has an equal chance to be a breakpoint (i.e., boundary of CNVs). However, this assumption is violated in the genetics perspective given the existence of inter-correlation among genomic positions, which is referred to as linkage disequilibrium (LD). Dictated by the presence of recombination hotspots that segment the genome into separate LD blocks, LD describes the correlated transmission of the alleles at adjacent locations in the genome. Interestingly, it was demonstrated early on that CNVs are outcomes of evolution and they originated from recombination-based processes [29]. These relationships demonstrate the possibility of the existence of CNV breakpoints located at the recombination hotspots, which violates the assumption of previous segmentation methods that assume each genetic location has an equal chance to be a CNV breakpoint. This further implies the importance of integrating the biological characteristics (i.e., LD structure) into statistical modeling for CNV detection.

Motivated by this fact, we here developed an accurate and fast segmentation algorithm by modeling the genomic correlation structure with a local search strategy for optimized computational efficiency. Early in 1996, Kim HJ. [30] explored a likelihood ratio test for non-independent observations, yet its computational complexity with its exhaustive search largely limited its application. To investigate the performance of the newly proposed algorithm, we conducted simulation studies to investigate its performance in single nucleotide polymorphs (SNP) array studies in a variety of scenarios. In this study, we demonstrated the improved performance of the novel algorithm in array-based real data analysis by using a set of “gold standard validation sets” of CNVs from the HapMap projects [31-33]. Overall, the new algorithm presented high sensitivity and accuracy in CNV detection, especially single copy changes. This new segmentation algorithm has a wide scope of application so it can be implemented in CNV detection tools for next-generation sequencing data analysis and single-cell sequencing analysis.

## 2. Materials and Methods

### 2.1 Notations and models

We use Y = (*Y*_1_, …, *Y*_*m*_)^*T*^ to denote the genetic intensities for a sequence with *m* biomarkers (e.g., SNPs in array data, or exon in whole exome sequencing data). For example, Y may present the log R Ratio (LRR) intensities of a chromosome from SNP array data. The model we consider is a change-point method with the basic model as

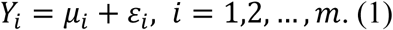

Its underlying mean *u* = (*u*_1_, …, *u*_*m*_)^*T*^ is a piecewise constant vector. A change-point method is to find change points defined as a position τ such that *μ*_τ_ ≠ *μ*_τ+1_. The locations of the change points therefore indicate the location of breakpoints or boundaries of CNVs. We assume that there are *M* change points in the sequence, 0 < τ_1_ < … < τ_*M*_ < *m*. Considering the sparsity of CNVs across the genome, we assume *m* is large and *M* is small. The goal is therefore to estimate the number of change points, *M*, and the location of the change points by the location vector, τ = (τ_1_, …, τ_*M*_)^*T*^. Many studies have worked on the problem of identifying the location of breakpoints when the *Y*’s are independent, such as CBS and screening and ranking algorithm (SaRa) [22, 26]. In this paper, to capture the biological characteristics in the process of copy number states inference, we assume the genetic intensities follow a multivariate normal distribution given the dependence structure of the genome (i.e., LD).

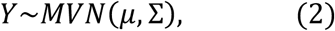

where Σ is the covariance matrix with dimension *m* × *m*. The covariance matrix (Σ) can be estimated by using the correlation matrix estimated from the samples or an exterior dataset such as samples from the 1000 Genomes project [34].

For a point *x*, a local diagnostic function *D*(*x*) is defined as the average mean difference in the observations before and after the point.

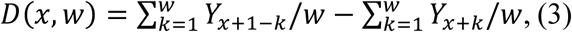

where *w* is the bandwidth. The quantity of *D*(*x, w*) depends on the local 2*w* data points 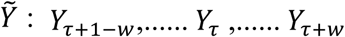 where 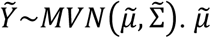 is a sub-vector of *μ* with length 2*w*; 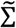 is a sub diagonal matrix of the covariance matrix Σ with dimension 2*w* × 2*w*, respectively. Then *D*(*x*) can be rewritten as 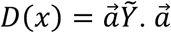 is a 2*w* vector takes the form 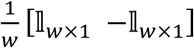. By derivation with the linear property of multivariate normal distribution, we obtained 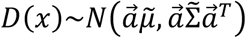. It turned out that the distribution of the local diagnostic function became a univariate normal with a covariance matrix depending on the local information, 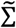. Since both 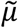 and 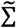 are known or can be estimated, the mean and variance of *D*(*x*) are functions of bandwidth *w* and only depend on the local sequence. As such, we proposed the algorithm, referred to LDcnv, based on a multivariate normal assumption that systematically integrated the biological characteristics into statistical modeling of the genetic intensities.

### 2.2 Copy number inference

After calculation of the local diagnostic statistic *D*(*x*), hypothesis testing is needed to find change-point candidates by a local screening and ranking strategy [26, 35]. A similar strategy has been used in our previous work [28] which guarantees computational speed of the CNV detection method in a whole genome scan.

Providing the distribution of *D*(*x*), we first define the *w*-local maximizer of a function. For any data point *x*, the interval (*x* − *w, x* + *w*) is called the *w*-neighborhood of *x*. And, *x* is a *w*-local maximizer of function *f*(·) if *f* reaches the maximum at *x* witin the *w*-neighborhood of *x*. In other words,

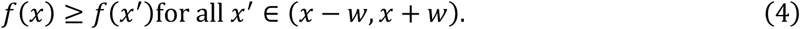

Then let ℒ be the set of all local maximizers of the function |*D* (*x, w*)| and we can select a subset 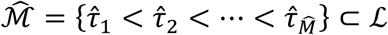 by setting a threshold 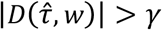, where 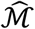 and 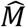 are the estimators for the locations and the number of change-points, respectively. To set up the threshold γ, we adopted a multiple comparison method using a false discovery rate approach (**Supplementary Text A1**). Empirical distribution of the local maximizers of the diagnostic statistic was generated mimicking the normal sequence with no change-points. As a result, the local maximizers 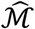 or, equivalently, local minimizers of p-values were selected.

Then we used the modified Bayesian Information Criteria (mBIC) to further eliminate false positives as proposed in [36]:

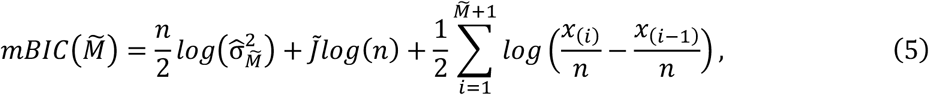

Where 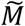 is all the possible values of 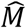 and 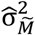 is the maximum likelihood estimator of the variance assuming 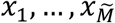 are change points. Then the final estimated number and the locations of change points are 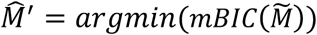 and 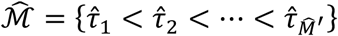, respectively. For copy number inference, Gaussian mixture model-based clustering was used for copy number state classification. Each segmented region will be classified using a five-state classification scheme (deletion of a single copy, deletion of double copies, normal/diploids, duplication of a single copy, and duplication of double copies) [37].

### 2.3 Numerical simulation studies

With the new proposed algorithm, we conducted extensive simulations to evaluate the performance in practice. We simulated SNP array data for demonstration of its advantage over existing algorithms.

To simulate the correlated genomic intensities, we used the first-order autoregressive (AR1) process:

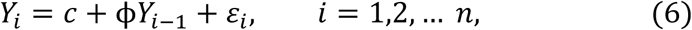

where *Y*_*i*_ was the intensities for the *i*-th marker; ε_*i*_ was a Gaussian white noise process with mean zero and variance 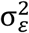; ϕ was a known coefficient that controlled the autocorrelation of the data series (for example, |ϕ| < 1 generates a stationary sequence); *c* was a constant and *n* was the total number of markers. The underlying mean *μ*, variance *var*(*Y*_*i*_) and auto-covariance *B*_*n*_ were given as: 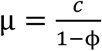, 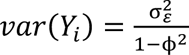 and 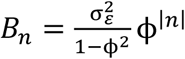. One advantage of using the AR1 process is that the change of the underlying distribution of the white noise term allows one to flexibly adjust the distribution of the data, especially when the normality is not satisfied. Also, with the AR1 process, we did not need to decompose the covariance matrix, which consequently generated data much faster than the multivariate normal distribution assumption-based process.

We randomly generated LRR and B Allele frequencies (BAF) intensities for 100 sequences (i.e., chromosomes) with 20,000 markers. For each sequence, 40 dispersed CNV segments were generated, the locations of which were randomly selected and were not overlapping with each other. The mean and variance were empirical values provided by the Illumina website (https://www.illumina.com/documents). We constructed different scenarios with different combinations of CNV sizes, status and correlation coefficient. The CNV sizes varied from 10∼50 markers, 50∼100 markers and 100∼200 markers. The CNV status included deletion of a single copy (Del.s), deletion of double copies (Del.d), duplication of a single copy (Dup.s) and duplication of double copies (Dup.d). To investigate the CNV data with different levels of correlations, the value of ϕ. which was equivalent to the Pearson’s correlation coefficient in theory, was set to be 0.1, 0.3 and 0.5, respectively. With the generated CNV data, we compared the proposed method to the performance of an independence assumption-based method, CBS, and a hidden Markov model based method, PennCNV [22, 38]. The PennCNV assumes a Markov chain; however, the LD structure is not directly incorporated in the statistical modeling. The performance of these methods was demonstrated by computing the true positive rate (TPR) and false positive rate (FPR).

### 2.4 Performance evaluation by application to HapMap datasets

To further assess the proposed LDcnv algorithm, we analyzed 180 healthy individuals with CNV profiles having been validated experimentally or statistically by three previous microarray studies [31-33]. The HapMap project utilized stringent genotyping quality control (QC) and merged results from multiple calling algorithms, which finally produced 856 high-quality CNV calls [33]. McCarroll et al. identified 1,320 high resolution CNV calls by joint analysis of multi-platforms data including Affymetrix SNP array, array CGH and fosmid end-sequence-pair, whereas Conrad et al. used tiling oligonucleotide array to generate a map of 11,700 CNVs, among which 8,599 were independently validated through stringent validation procedures such as quantitative PCR [31, 32]. The SNP array data were downloaded from the international HapMap 3 Consortium [33]. All individuals were Utah residents with Northern and Western European ancestry (CEU). Genotype data were generated by the genotyping platform Affymetrix Human SNP array 6.0. Specifically, a stringent QC procedure was adopted (e.g., the CNV must overlap with 2 to 20 exons with less than 5% missing rate across all samples) to generate high-quality CNV profiles.

Using the final “gold standard validation sets”, we compared the performance of the LDcnv method against PennCNV and CBS [22, 38]. To obtain high-quality CNV profiles, we excluded CNVs with less than ten markers in the calling results. Besides, we used the database of genomic variants (DGV) [17] as a reference of common variants as a quality control step to keep the high-quality CNV profile. DGV curates CNV records from 55 independent studies of clinically normal populations with 202,431 CNV regions.

These methods were assessed by the precision rate, recall rate and F1 score measures. The precision rate was defined as the ratio of identified true positives over the total number of identified CNVs. The recall rate was the ratio of identified true positives over the total number of “true CNVs” in the “gold standard validation sets”. The F1 score was defined as the harmonic mean of precision and recall rate which reflected the overall accuracy. Moreover, we also evaluated the performance in subsets of the validation sets of those that were less than ten markers to assess the performance in detecting short CNVs.

### 2.5 Theoretical Derivation: Correlation Structure in CNV data

In this study, we hypothesized that the integration of biological characteristics will increase the accuracy of CNV detection. In this section, we therefore provide theoretical evidence to support that the genetic intensity data are presenting a correlation structure that should be deliberated in statistical modeling for CNV detection.

We start from the generation of the two SNP array intensities, LRR and BAF, which has been introduced in Wang et al. [38] and summarized here. For one SNP with two alleles defined as *A* and *a*, the raw signal intensity values are measured for each allele and then are processed with a five-step normalization procedure using the information of all SNPs. *X* and *Y* values are produced for each SNP, representing the normalized signal intensity on the *A* and *a* alleles, respectively. Two additional measures are then calculated for each SNP, where *R* = *X* + *Y* representing the total signal intensity, and 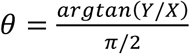 referring to the relative allelic signal intensity ratio. As a normalized measure of total signal intensity, LRR measures the normalized total intensity of the possible alleles for a given marker, from which the magnitude of mean changes (or called jump size) are used for inference of the boundaries of CNVs (i.e., breakpoints). The LRR value for each SNP is calculated as *LRR* = *log*_2_(*R*_*observed*_/*R*_*expected*_) where *R*_*expected*_ is computed from linear interpolation of canonical genotype clusters [39].

In our study, to present the correlation of two adjacent bi-allelic SNPs, we assume the reference allele and alternative allele were *A* and *a* for the first SNP, and *B* and *b* alleles for the second SNP. The total signal intensities for the two alleles are therefore *X*_*A*_ + *Y*_*A*_ and *X*_*B*_ + *Y*_*B*_. We assume that we observe the two reference alleles *A* and *B* with frequencies *p*_*A*_ and *p*_*B*_ in the whole population. Under the Hardy-Weinberg equilibrium assumption, the joint probability for the nine genotypes can be calculated (shown in Table 1). For example, for the genotype *AABB*, the genotype frequency will be (*p*_*A*_*p*_*B*_ + *D*_*AB*_)^2^ where *D*_*AB*_ is the coefficient of linkage disequilibrium between the two SNPs.

**Table 1:** Joint genotype probabilities for two diallelic loci. The joint genotype probabilities were calculated under the Hardy-Weinberg equilibrium assumption. *A* and *a* are reference and alternate alleles in the first locus, *pA* is the probability of the reference allele; *B* and *b* are the reference and alternate alleles in the second locus, *pB* is the probability of the reference allele; *D*_*AB*_ is the coefficient of linkage disequilibrium between two loci.

| Locus 1 | Locus 2 | Probabilities |
| --- | --- | --- |
| AA | BB | $(p_A p_B + D_{AB})^2$ |
| AA | Bb | $2(p_A p_B + D_{AB})(p_A(1 - p_B) - D_{AB})$ |
| AA | bb | $(p_A(1 - p_B) - D_{AB})^2$ |
| Aa | BB | $2(p_A(1 - p_B) - D_{AB})((1 - p_A)p_B - D_{AB})$ |
| Aa | Bb | $2(p_A p_B + D_{AB})((1 - p_A)(1 - p_B) + D_{AB}) + 2(p_A(1 - p_B) - D_{AB})((1 - p_A)p_B - D_{AB})$ |
| Aa | bb | $2(p_A(1 - p_B) - D_{AB})((1 - p_A)(1 - p_B) + D_{AB})$ |
| aa | BB | $((1 - p_A)p_B - D_{AB})^2$ |
| aa | Bb | $2((1 - p_A)p_B - D_{AB})((1 - p_A)(1 - p_B) + D_{AB})$ |
| aa | bb | $((1 - p_A)(1 - p_B) + D_{AB})^2$ |

First, to calculate the correlation of LRR between the two SNPs, we need to compute the correlation of the non-linear logarithm transformation of the *observed* total signal intensities *log*.(*R*_*observed*_/*R*_*expected*_). The *observed* total signal intensities, *R*_*observed*_, can be calculated from the dataset, and the value of *R*_*expected*_ is a fixed value. After applying the Taylor expansions [40], the correlation of the LRR intensities can be approximately represented by the correlation of *R*_*observed,A*_ and *R*_*observed,B*_, which is expressed by

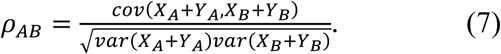

Derivation of the *p*_*AB*_ will show the correlation structure of the LRR intensities, which will be further discussed in the results **(Section 3.1)**.

## 3. Results

### 3.1 Theoretical proof revealed the correlation structure in intensity signals

First, to further demonstrate the necessity of integrating the correlation structure into the segmentation algorithm, we initiated a theoretical derivation to demonstrate the correlation structure of the genetic intensities (i.e., LRR) between two adjacent genomic loci. As a continued discussion of Section 2.5, we have the correlation coefficient of LRR between two loci expressed as 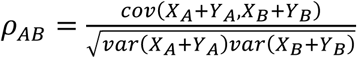, in which *X* and *Y* are the normalized signal intensities of the two alleles in a SNP (e.g., *A* and *a*).

For *cov*(*X*_*A*_ + *Y*_*A*_, *X*_*B*_ + *Y*_*B*_), we obtained

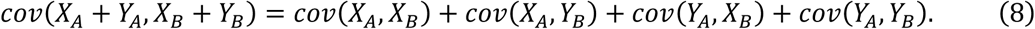

As *cov*(*X*_*A*_, *X*_*B*_) = *E*(*X*_*A*_*X*_*B*_) − *E*(*X*_*A*_)*E*(*X*_*B*_), the *expected* values of the normalized signal intensities *X*_*A*_, *X*_*B*_ and the *expected* values of their product need to be derived. We assume the joint probability density function to be 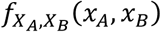 which are bivariate normal distributions conditional on the genotype *G*:

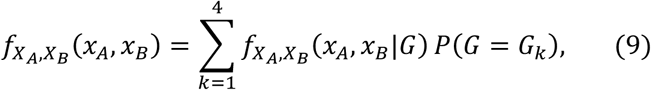

where *G* = [*AABB, AABb, AaBB, AaBb*]^*T*^ is the vector of genotypes that contain alleles *A* and *B*. After mathematical derivation (detailed in **Supplementary Text A2**), the covariance between the two normalized signal intensities can be formulated as:

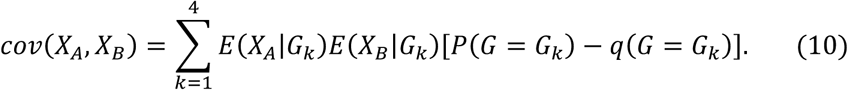

*q*(*G* = *G*_*k*_) is the genotype frequency under the condition that the two loci are in LD. For example, 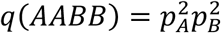. The expression of all the other genotype frequencies *P*(*G* = *G*_*k*_) can be found in Table 1.

Similarly, we can derive the other three terms in equation (8) and then obtain *cov*(*X*_*A*_ + *Y*_*A*_, *X*_*B*_ + *Y*_*B*_) as

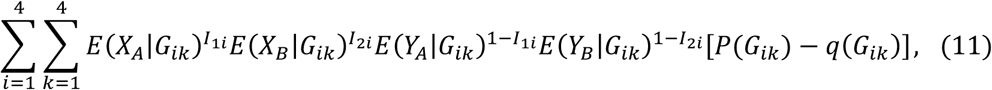

where *G*_1._ = [*AABB, AABb, AaBB, AaBb*]^*T*^, *G*_2._ = [*AABb, AAbb, AaBb, Aabb*]^*T*^, *G*_3._ = [*AaBB, AaBB, aaBB, aaBb*]^*T*^ and *G*_O._ = [*AaBb, Aabb, aaBb, aabb*]^*T*^.*I*_*1i*_ and*I*_2*i*_are indicator functions of whether *X*_*A*_ and *X*_*B*_ contribute to the bivariate density in equation (9).

*I*_1*i*_ = 1, if *i* = 1 or *I*_2*i*_ = 1, if *i* = 1 or 3. Otherwise, *I*_1*i*_ = *I*_2*i*_ = 0.

Combining results from equation (11) and the expression of the denominator of *p*_*AB*_ in equation (7) from **Supplementary Text A3**, the correlation of LRR between the two loci can be defined as:

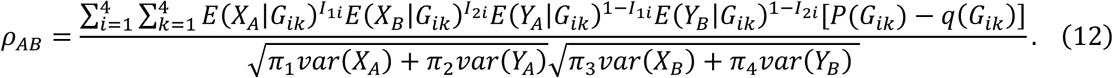

According to expression of Equation (12), the correlation of LRR depends on the correlation of the two SNPs which was measured by the LD coefficient *D*_*AB*_. For example, 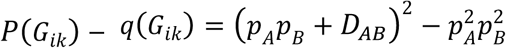 for genotype *AABB* (*i* = 1, *k* = 1). *p*_*AB*_= 0 if D_*AB*_= 0.

In summary of this section, we found that the correlation of LRR between two loci are related to the coefficient of LD measure, although the relationship does not admit a simple format.

### 3.2 Real data shows that that CNV locations are related to the Genomic Structure

To explore the relationship between CNV locations and LD structure, we utilized the high-quality CNV profile from the international HapMap 3 Consortium which merged probe-intensity data from both Affymetrix and Illumina arrays [33]. The CNV profile set contained 856 records for 1,184 individuals. We randomly selected 300 high-quality CNVs and mapped them to the LD block maps (Figure 1). Obviously, most of these CNVs are located outside of the LD blocks (across block, hybrid or random), with only 2.0% residing within LD blocks (inter-block) (Supplementary Table 1). Among the CNV types that do not involve LD structure (i.e., across block, inter-block and hybrid), there were only 10.7% of the CNVs were located within blocks. These results implicated that CNVs are not randomly distributed across the genome, and their distribution is closely related to the local LD structure. Such results motivated the development of the LDcnv algorithm and are consistent with the theory derivation of the correlations structure in SNP array data (**Section 3.1**).

**Figure 1.**
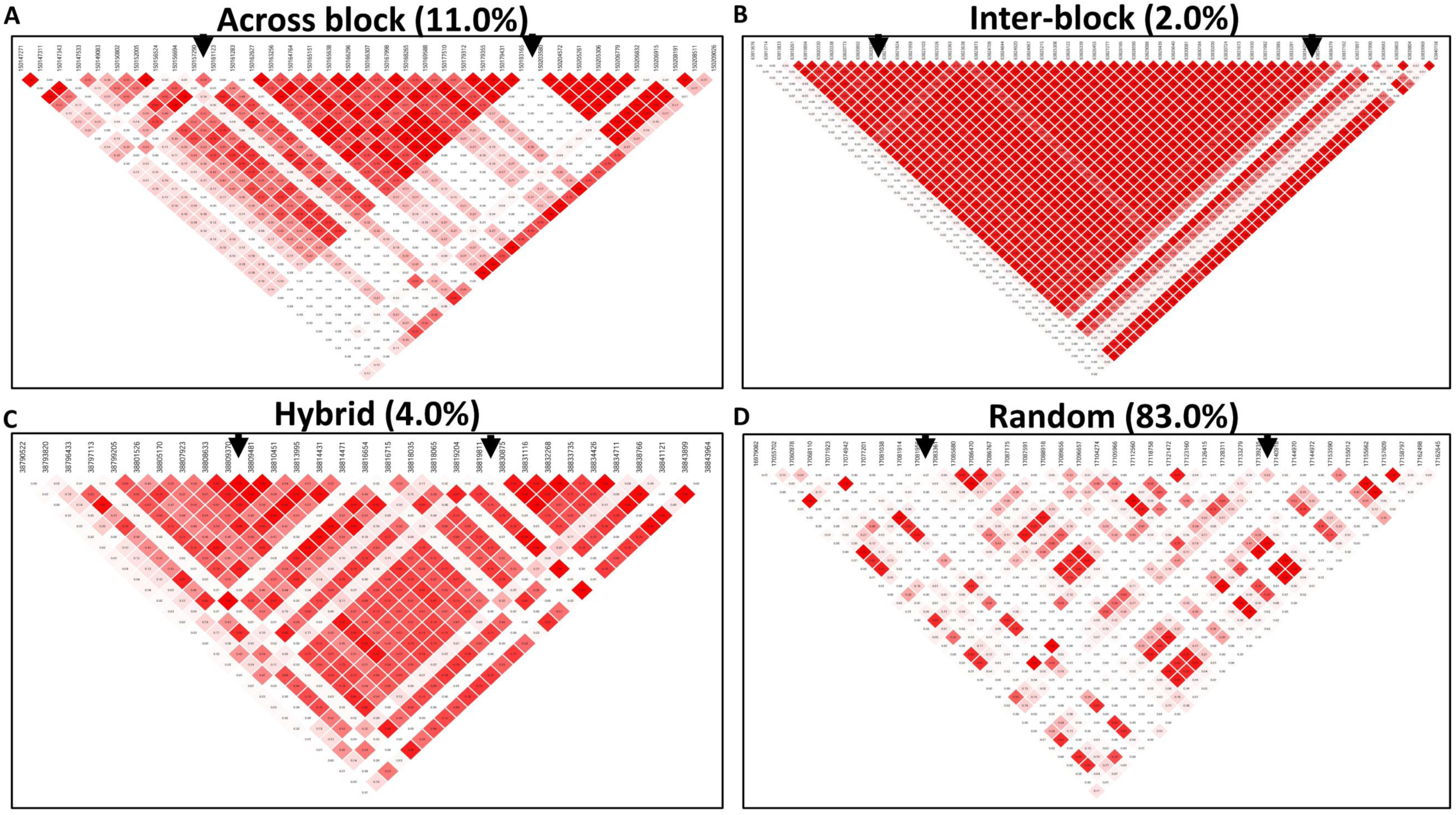
Four classifications of the CNV locations in the LD genome map. The graphs summarized the frequency of CNV types with the existing high-quality CNVs from the HapMap phase 3 project. (a) Across block: CNVs spanning at least one LD blocks, (b) Inter-block: CNVs locate within a LD block, (c) Hybrid: only one breakpoint locating within LD block, and (d) Random: CNVs locating in the area with weak or no LD structure. The black arrows in each plot note the start and end points of the CNV.

### 3.3 Simulation studies show improved performance of LDcnv

First, we used the simulated data to evaluate the performance of the LDcnv method in SNP array analysis under a variety of scenarios: (1) different correlation level; (2) different CNV sizes; and (3) different CNV status (see Methods). For data with moderate correlation coefficient (ϕ = 0.3) that is assumed to be close to real data, the LDcnv method presented a consistent power gain in detecting CNVs from single copy duplication/deletions (i.e., Dup.s and Del.s) to double copy changes (i.e., Dup.d and Del.d) (Table 2). The performance of the LDcnv method was obviously superior to the other two methods when the CNVs had small jump sizes (Dup.s and Del.s), especially for short CNVs (<200 markers). For example, when the CNVs had a length between 10-50 markers and the CNV status was single copy duplication (Dup.s), the LDcnv method had a TPR at 0.88, while the FPR was 0.12. The corresponding TPRs and FPRs for PennCNV were 0.84 and 0.11; and 0.69, 0.32 for CBS, respectively. When the CNV size increased from 10-50 markers to 100-200 markers, the LDcnv methods presented stable estimations of CNVs, whereas the other two methods had diminished power. A similar pattern was *observed* when the correlation increased from 0.1 to 0.5 (Table 3).

**Table 2:** Summary of CNV calls on simulated data at ϕ =0.3 from all methods. True positive rates (TPRs) and false positive rates (FPRs) of LDcnv, PennCNV and CBS with different CNV states and CNV sizes, the autoregressive coefficient (ϕ) was fixed at ϕ =0.3 which was corresponding to Pearson’s correlation coefficient at 0.3. Del.d: deletion of double copies; Del.s: deletion of single copy; Dup.s: duplication of single copy; Dup.d: duplication of double copies.

| CNV State | Method | CNV length (markers) |  |  |  |  |  |
| --- | --- | --- | --- | --- | --- | --- | --- |
|  |  | 10~50 |  | 50~100 |  | 100~200 |  |
|  |  | TPR | FPR | TPR | FPR | TPR | FPR |
| Del.d | LDcnv | 0.99 | <0.01 | 0.97 | <0.01 | 0.99 | 0.01 |
|  | PennCNV | 1.00 | <0.01 | 1.00 | <0.01 | 1.00 | <0.01 |
|  | CBS | 1.00 | 0.04 | 1.00 | 0.07 | 1.00 | 0.09 |
| Del.s | LDcnv | 0.99 | 0.02 | 0.99 | 0.02 | 0.99 | 0.03 |
|  | PennCNV | 0.96 | 0.03 | 0.95 | 0.04 | 0.86 | 0.11 |
|  | CBS | 0.98 | 0.03 | 0.98 | 0.04 | 0.98 | 0.05 |
| Dup.s | LDcnv | 0.97 | 0.03 | 0.94 | 0.03 | 0.93 | 0.05 |
|  | PennCNV | 0.91 | 0.07 | 0.92 | 0.08 | 0.87 | 0.11 |
|  | CBS | 0.85 | 0.11 | 0.88 | 0.12 | 0.88 | 0.13 |
| Dup.d | LDcnv | 1.00 | <0.01 | 1.00 | <0.01 | 0.99 | <0.01 |
|  | PennCNV | 1.00 | <0.01 | 0.99 | <0.01 | 0.99 | 0.01 |
|  | CBS | 1.00 | 0.01 | 1.00 | 0.02 | 1.00 | 0.04 |

**Table 3:** Summary of CNV calls on simulated data at ϕ =0.5 from all methods. True positive rates (TPRs) and false positive rates (FPRs) of LDcnv, PennCNV and CBS with different CNV states and CNV sizes, the autoregressive coefficient (ϕ) was fixed at ϕ =0.5 which was corresponding to Pearson’s correlation coefficient at 0.5. Del.d: deletion of double copies; Del.s: deletion of single copy; Dup.s: duplication of single copy; Dup.d: duplication of double copies.

| CNV State | Method | CNV length (markers) |  |  |  |  |  |
| --- | --- | --- | --- | --- | --- | --- | --- |
|  |  | 10~50 |  | 50~100 |  | 100~200 |  |
|  |  | TPR | FPR | TPR | FPR | TPR | FPR |
| Del.d | LDcnv | 0.99 | 0.01 | 0.96 | 0.01 | 0.99 | 0.03 |
|  | PennCNV | 0.99 | 0.01 | 0.99 | 0.01 | 1.00 | <0.01 |
|  | CBS | 1.00 | 0.23 | 1.00 | 0.44 | 1.00 | 0.64 |
| Del.s | LDcnv | 0.96 | 0.06 | 0.95 | 0.08 | 0.96 | 0.09 |
|  | PennCNV | 0.89 | 0.05 | 0.88 | 0.09 | 0.83 | 0.14 |
|  | CBS | 0.94 | 0.26 | 0.94 | 0.46 | 0.95 | 0.62 |
| Dup.s | LDcnv | 0.88 | 0.12 | 0.89 | 0.09 | 0.91 | 0.11 |
|  | PennCNV | 0.84 | 0.11 | 0.86 | 0.13 | 0.83 | 0.18 |
|  | CBS | 0.69 | 0.32 | 0.80 | 0.57 | 0.79 | 0.76 |
| Dup.d | LDcnv | 0.99 | 0.01 | 0.96 | 0.01 | 0.99 | 0.03 |
|  | PennCNV | 0.99 | 0.01 | 0.99 | 0.01 | 1.00 | <0.01 |
|  | CBS | 1.00 | 0.23 | 1.00 | 0.44 | 1.00 | 0.64 |

In conclusion, the LDcnv method which integrated the correlation structure in the model, largely presented overall high accuracy, stability and robustness in CNV detection, especially for detection of CNVs with small jump sizes.

### 3.4 Application to the HapMap datasets

We further applied the LDcnv method to a real data study in comparison with CBS and PennCNV [22, 38]. Using the DGV as a common variant reference database, 79.65% of the CNVs identified by LDcnv have been reported as common variants that are not diseases relevant which implicated the validity of the CNV calls from our method.

With the validation sets from the three datasets (including HapMap3, Conrad et al., and McCarrol et al.), the total number of “true” CNVs included in the three sets were 19,936, 121,453 and 11,961, separately. Among them, 10,005, 98,387 and 5,277 were short CNVs with length less than ten markers. The overall accuracy of the LDcnv method was greater than that of the two other methods in all three validation sets (Table 4, Figure 2). Specifically, LDcnv presented higher recall rate or sensitivity that it detected more true positives. For detection of short CNVs (Table 5, Figure 3), the LDcnv method was comparable to the other methods in all three validation sets and it presented obviously higher precision or specificity in detecting short CNVs. It is noteworthy that CBS was the most sensitive one that detected the highest number of variants which was consistent with previous findings showing that CBS was good at calling the exact boundaries of CNVs [44], but such high recall rate came at the expense of precision. CBS also has the lowest computational speed among these three methods. As *expected*, PennCNV was the most conservative one that detected the lowest number therefore presented the lowest prevision rate (Table 5). These results further demonstrated that the integration of correlation structure significantly improved the overall performance of CNV detection.

**Table 4:** Overall assessment of CNV calling on the HapMap project dataset. Performance assessment of CNV calls from the HapMap Project 3 in the 180 HapMap samples by LDcnv, PennCNV and CBS on reports from (a) HapMap3 (b) Conrad et al. (c) McCarroll (MCC) et al. studies. The recall rate was defined as the ratio of identified true positives over the total number of “true CNVs”. The F1 score was calculated as harmonic mean of precision rate and recall rate. TP: True positives among the detected CNVs.

|  | HapMap3 |  |  |  | Conrad |  |  |  | MCC |  |  |  |
| --- | --- | --- | --- | --- | --- | --- | --- | --- | --- | --- | --- | --- |
|  | TP | Precision | Recall | F1 | TP | Precision | Recall | F1 | TP | Precision | Recall | F1 |
| LDcnv | 4463 | 52.72% | 22.30% | 31.42 | 5888 | 64.99% | 4.84% | 9.02 | 2861 | 63.53% | 23.91% | 34.75 |
| PennCNV | 3760 | 53.23% | 18.81% | 27.85 | 4880 | 64.56% | 4.01% | 7.56 | 2640 | 66.48% | 22.07% | 33.14 |
| CBS | 4044 | 55.41% | 20.18% | 29.69 | 5099 | 65.02% | 4.19% | 7.88 | 2572 | 64.94% | 21.40% | 32.30 |

**Table 5:** Assessment of calling performance in short CNVs on the HapMap project dataset. Performance assessment on detecting short CNVs (<10 markers) from the HapMap Project 3 in the 180 HapMap samples by LDcnv, PennCNV and CBS on reports from (a) HapMap3 (b) Conrad et al. (c) McCarroll (MCC) et al. studies. The recall rate was defined as the ratio of identified true positives over the total number of “true CNVs”. The F1 score was calculated as harmonic mean of precision rate and recall rate. TP: True positives among the detected CNVs.

|  | HapMap3 |  |  |  | Conrad |  |  |  | MCC |  |  |  |
| --- | --- | --- | --- | --- | --- | --- | --- | --- | --- | --- | --- | --- |
|  | TP | Precision | Recall | F1 | TP | Precision | Recall | F1 | TP | Precision | Recall | F1 |
| LDcnv | 963 | 10.40% | 9.62% | 10.00 | 1757 | 16.39% | 1.78% | 3.22 | 703 | 14.16% | 13.32% | 13.72 |
| PennCNV | 177 | 2.65% | 1.76% | 2.12 | 698 | 8.78% | 0.70% | 1.31 | 206 | 5.33% | 3.90% | 4.51 |
| CBS | 1279 | 8.48% | 12.78% | 10.19 | 2933 | 16.63% | 2.98% | 5.05 | 850 | 10.59% | 16.10% | 12.78 |

**Figure 2.**
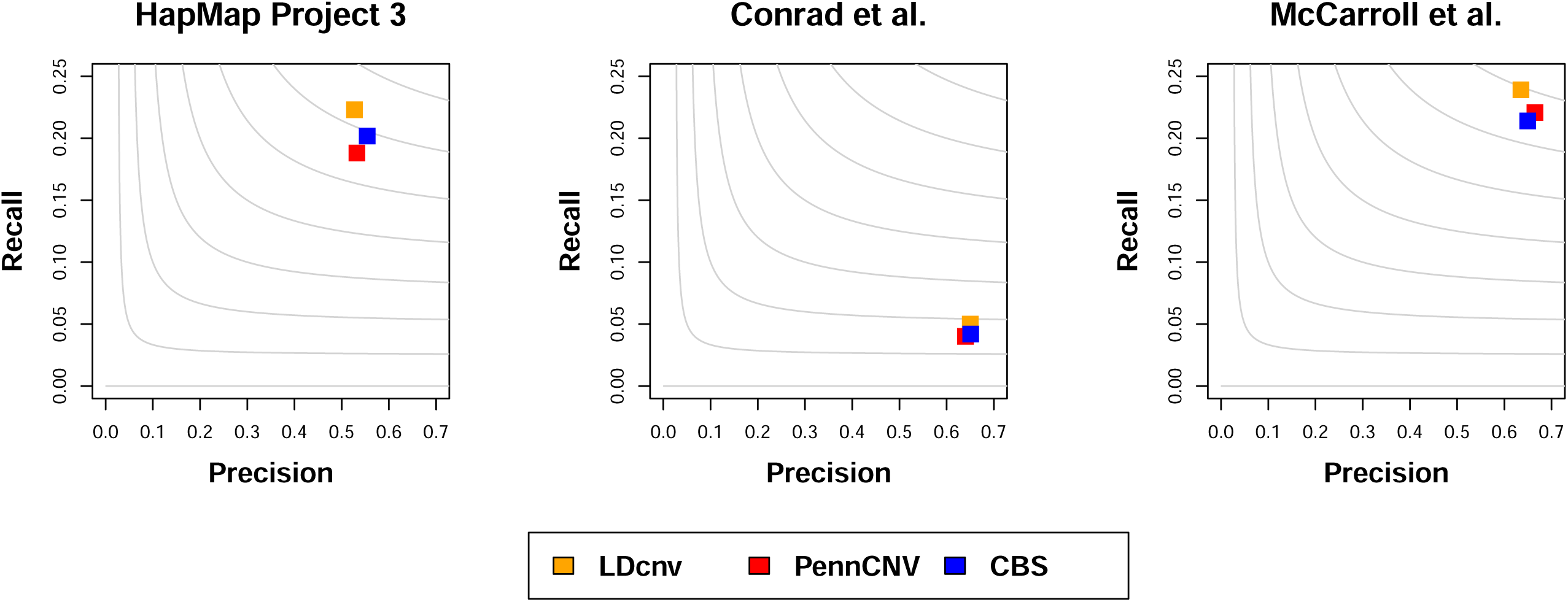
Assessment of CNV calls generated by LDcnv, PennCNV and CBS with validation CNV calls. Performance of the LDcnv, PennCNV and CBS methods in detection validated CNVs from (a) HapMap 3 (b) Conrad et al (c) McCarroll et al. The grey contours are F1 scores calculated as the harmonic mean of precision rate and recall rate.

**Figure 3.**
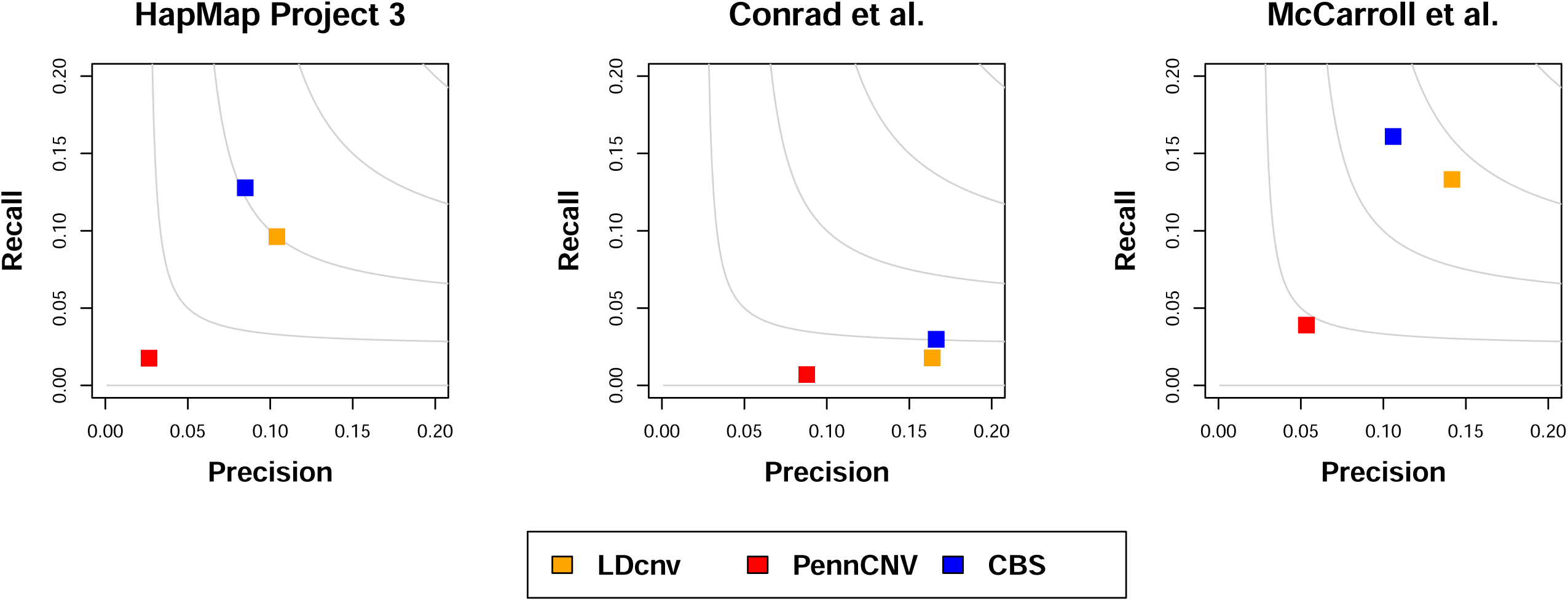
Assessment of CNV calls calling performance in short CNVs on the HapMap project dataset. Performance assessment on detecting short CNVs (<10 markers) from the HapMap Project 3 in the 180 HapMap samples by LDcnv, PennCNV and CBS on reports from (a) HapMap3 (b) Conrad et al. (c) McCarroll (MCC) et al. studies. The grey contours are F1 scores calculated as the harmonic mean of precision rate and recall rate.

## 4. Discussion

CNVs play important roles in human complex diseases [1-7].While numerical CNV detection tools have been developed for modern genotyping technologies, current detection methods are using segmentation algorithms that mainly focus on detecting random signals and do not fully address the CNV-specific challenges. These challenges include the multiple natural features of a CNV, including dependent random noise signals and small jump sizes of the breakpoints. In this work, we introduce a correlation-based segmentation algorithm for CNV detection analysis that accommodates the non-independence nature of the genetic intensities. Simulation and real data analyses suggested that the LDcnv algorithm presented stable performance across different scenarios of CNV sizes, states and correlation coefficients, and it had a better or comparable accuracy compared to the independence assumption methods. The largest power gain tended to occur when CNVs were short and with small jump sizes, e.g., the duplication of a single copy.

This is the first report that demonstrates the promise of improved CNV detection by integrating the biological characteristics (i.e., LD structure) into statistical modeling. Utilization of these biological characteristics of CNVs improved accuracy and boosted power with the noisy and complex data. Instead of assuming equal weight on each chromosomal site, the LDcnv algorithm tends to put more weight on recombination hotspots which are more likely to be CNVs. For example, those CNVs with specific LD block structures missed by traditional algorithms will be identified by the new algorithms. This approach will also provide a valuable knowledgebase for CNV detection methodology development to integrate the genomic structure that delivers comprehensive information among genes.

In this study, we first presented the theoretical derivation of the correlation structure of the genetic intensity data from SNP array data. Well-developed array-based CNV analytical tools are usually based on segmentation and smoothing of LRR and BAF [38]. We found that the array-based LD structure, which was computed from the genotype frequencies, can be reflected in the correlation structure of the genetic intensity data. We stated that the correlation between two loci in the genetic intensity will depend on the LD coefficient computed from SNP allele frequencies. This evidence provides strong evidence of the existence of genomic correlation structure in the CNV data.

By implementing the correlation structure in the statistical model, we showed that the LDcnv algorithm presented essential advantages over the other independence assumption methods (e.g., CBS and PennCNV). These advantages are demonstrated by the simulation studies and the HapMap project with the “gold standard validation sets”. The superiority of the LDcnv algorithm over PennCNV was further demonstrated, especially in detecting short CNVs, which is the most difficult copy number states to be detected due to the embedded undetectable signal in the random noises. A possible explanation for this phenomenon is that short CNVs tend to have more evident correlation structure when they are located within an LD block. Such a characteristic cannot be easily captured by the hidden Markov model adopted in PennCNV, which assumes a constant level of dependence across the genome.

Indeed, the clinical relevance of small CNVs has been demonstrated in many studies in recently years. For example, Reza et al. [41] investigated a cohort of 714 patients with neurodevelopment disorders and verified the diagnostic importance of small CNVs. However, due to the noise of genotyping data, small segments are usually very difficult to distinguish from the normal noise signals. As such, the LDcnv algorithm may serve as an important tool for detecting small CNVs. Our result in simulations and the analyses of the HapMap CEU samples showed that LDcnv was capable of capturing those signals.

With LDcnv, we address the non-independence noise signal assumption by introducing a covariance matrix in the statistical modeling. To retain the covariance structure in the model, we can either use the correlation matrix estimated from the samples in the data or LD-based computation from reference samples (i.e., samples from the 1000 Genomes project). The advantage of the data-based estimation of the covariance relies on its feasibility and simplicity; however, the covariance might be data specific and the computational concerns will be encountered for large sample sizes. In contrast, the LD-based estimates with information coming from the population level might be more stable but be susceptible to a specific population substructure. As discussed in Mathew et al. [42], an alternative way is to use the map functions (e.g., the Haldane function) in an exponential function to estimate the covariance structure on each chromosome.

Moreover, the current study of the LDcnv algorithm is mainly focuses on its application to SNP array data, but it has great potential to be implemented in the whole exome sequencing (WES) or whole genome sequencing (WGS) data analysis. The main challenges for sequencing data come from the high level of biases and artifacts effects, which therefore require a well-designed normalization procedure. Another difficulty is that the WES data are count data that requires discrete variable distribution. It has been suggested that read counts are not appropriately modeled by the normal assumption model, even after a commonly used log-based transformation is applied [43]. To allow the implementation of the LDcnv algorithm, the modelling framework and segmentation procedure need to be adjusted accordingly in NGS data analysis.

Through theoretical derivation, it is the first report in which we showed the relationship between the SNP-based LD coefficient and the correlation coefficient in the intensity data. We have demonstrated that the LD structure can be reflected in the genetic intensity data (i.e., LRR). However, the correlation structure of the other important source of information, the BAF intensities, was not clear and not easily constructed. The theoretical study of BAF and the implementation of this information requires future studies. In addition, we used 300 high-quality CNV data from HapMap to demonstrate that CNVs were not randomly distributed across the genome. One possible explanation might be that the LD blocks under study were not long enough to contain a CNV. Rigorous examinations of this assumption with shorter CNVs should be further studied in the future. Still, the LDcnv algorithm may open a door for integrating biological characteristics in CNV detection methodology development with change-point methods.

## Supporting Information

**Supplementary Text A1.** This section includes the detailed information about the FDR approach that is used in the LDcnv algorithm.

**Supplementary Text A2-A3.** This section includes the detailed derivation of components in the correlation structure in CNV data.

**Supplementary Tables.** The supplementary tables are attached to support the study are attached.

## Acknowledgements

We thank the reviewers in advance for their thoughtful and insightful comments.

## Author Contributions

The conception and design of this study were developed by FX. The theoretical derivation and methodology development were conducted by XL and FX. The data analysis and interpretation were conducted by XL, FX, CG and FQ. The writing, review and manuscript revision were conducted by FX, XL and CG.

